# Loop engineering and activity improvement of TEV protease by a phagemid-based selection system

**DOI:** 10.64898/2025.12.03.692113

**Authors:** Hiroki Yamaguchi, Mark Isalan

**Author notes:** Corresponding authors: Mark Isalan, Department of Life Sciences, Imperial College London, Exhibition Road, SW7 2AZ London, UK.

## Abstract

Loop engineering of enzymes remains challenging due to high flexibility and conformational complexity, posing a bottleneck for deep-learning-based design. Here, we constructed mutant libraries for three loops of TEV protease to assess combining directed evolution with deep learning. Using an M13 phagemid-based selection system, the three libraries were screened, resulting in a Loop 1 variant (HyperTEV60/L1) that significantly enhanced the Michaelis constant (*K*_m_) of the HyperTEV60 scaffold, a highly active mutant identified by ProteinMPNN. Structural modeling suggested that a single-residue deletion and substitution in Loop 1 expands the substrate binding pocket, accounting for the improved *K*_m_. Although the catalytic efficiency *k*_cat_/*K*_m_ of HyperTEV60/L1 was only marginally higher than HyperTEV60, due to a *k*_cat_ decrease, our results reveal that the phagemid-based selection system tended to find variants optimizing *K*_m_. This study demonstrates that combining deep-learning-based global optimization with localized directed evolution maximizes the probability of discovering distinct, high-performance enzyme variants.

## Introduction

Proteins are fundamental components of life, operating as enzymes, structural components, and signaling molecules^1, 2, 3, 4, 5^. Through the elucidation of the intricate structural and functional properties of proteins, researchers have achieved a more profound comprehension of living systems. Moreover, the development of proteins for industrial use has been pursued through the design and screening of proteins with specific functions. In the early days of industrial enzyme development, readily available wild-type enzymes were primarily utilized for applications such as food processing, laundry detergents, and materials production^6, 7^. Subsequent advancements in molecular biology technologies have facilitated the re-engineering of existing enzymes in nature, resulting in variants that exhibit enhanced properties in terms of activity, specificity, and stability^8^. In recent years, research has also been active in *de novo* design to create novel sequences with desired properties and in conferring catalytic activity to protein scaffolds^9^. While such approaches are still in their infancy, the efficient exploration of the vast sequence space has proven to be a significant challenge. Nonetheless, it is anticipated that the application of this technology to industrial proteins will undergo expansion in the near future. In order to surmount the challenges, deep-learning-based tools for protein sequence design are rapidly gaining traction and have successfully generated novel proteins exhibiting excellent expression, and solubility^10, 11, 12, 13^.

ProteinMPNN, a deep-learning-based tool, has been demonstrated to generate highly stable sequences for a designed structural backbone^14^. In addition, for native structures, it has been shown to generate sequences that are predicted to fold reliably into the intended structure, sometimes exhibiting enhanced properties compared to the native sequence^14, 15^. The development of this computational tool was driven by the necessity to reproduce the benefits of experimental methods such as directed evolution, while concomitantly minimizing the requirement for extensive resources and labor. Indeed, as K. Sumida *et al*. demonstrate in their study, a strategy was developed to enhance solubility and stability by applying ProteinMPNN to natural protein sequences^15^. Utilizing TEV protease as a model, they designed and evaluated multiple novel variants, identifying mutants that exhibited significant enhancement in solubility, yield, and thermal stability, as well as in catalytic efficiency (*k*_cat_ / *K*_m_). Notably, their analysis via molecular dynamics (MD) simulations suggested that changes in the flexibility of specific loop regions contributed to this enhanced activity.

It is widely acknowledged within the scientific community that loop regions are of particular importance in the field of enzyme engineering^16^. This is due to the fact that these regions are often involved in the overall conformational changes and regulation of catalytic activity in the three-dimensional structure of enzymes^17, 18, 19^. Indeed, the pivotal function of loops in substrate selectivity, specificity, and activity has been documented in numerous enzymes^20, 21, 22, 23, 24^. However, loop regions are generally highly dynamic, often becoming disordered in X-ray crystallography or Cryo-EM single-particle analysis, resulting in a paucity of experimental structural information. Furthermore, their involvement in multiple conformational changes that are essential for catalysis makes the construction of clear loop function databases difficult, which poses a significant bottleneck for the design of deep-learning-based methods targeting loop engineering^25^.

The fundamental question this raises is therefore as follows: can existing deep-learning-based methods adequately cover the entire search space for proteins when engineering loops that influence enzyme activity and specificity, or are further improvements possible? The objective of this study is therefore to gain insight into the enzyme optimization landscape by conducting directed evolution in specific regions of the ProteinMPNN-optimized TEV protease model and examining the impact of these loops on activity. The findings of this study demonstrate the potential for directed evolution to discover sequences distinct from those found by deep-learning-based methods. This suggests an effective development strategy for industrial enzyme creation. Ultimately, we propose the combination of global optimization by ProteinMPNN with local optimization by directed evolution, thereby re-emphasizing the complementary value of exploring the experimental search space for loop sequence optimization.

## Results and discussion

### Library design of TEV protease

In 2024, K. Sumida, *et al.* reported an improvement in the functions of TEV protease using ProteinMPNN^15^. In that study, an autolysis-resistant S219D variant^26^ (henceforth designated TEVd) was utilized as a template. Focusing on the sequence conservation within the TEV protease family, the authors created a total of four datasets: one preserving only the active site residues, and three others preserving the active site residues plus residues with 30%, 50%, and 70% identity, respectively. Subsequently, 144 sequences were designed from these datasets using ProteinMPNN. The designed sequences were evaluated using AlphaFold2 to determine whether they formed the same structure as TEVd, thereby confirming that their predicted Local Distance Difference Test (pLDDT) scores were ≥ 87.5 (the native TEV is predicted with pLDDT = 90). Following a screening process, a variant was identified that was termed "hyperTEV60". This variant demonstrated an approximate 26-fold increase in *k*_cat_ /*K*_m_ in comparison to TEVd. The results of the MD simulations indicated that the activity in question was the result of the contribution of three loops.

The present study concentrated its investigation on HyperTEV60, the variant that exhibited the most significant activity enhancement^15^. Initially, a comparison was made between the sequences of the template, TEVd, and HyperTEV60 (Fig. S1). The three loops that have been hypothesized to be responsible for the increased activity were observed to differ by 4, 5, and 2 residues between the two sequences, respectively (see Fig. 1 and S1). In order to investigate the effects of these three loops (Loop 1, 2, and 3) on TEV protease activity, mutant libraries were initially constructed for each loop. For the residues that differed between TEVd and the ProteinMPNN-designed HyperTEV60, combinatorial libraries were generated by randomizing each position using NNK codon primers to minimize stop codons (where N = A, C, G or T and K = G, T). In principle, given the capacity for each position to be substituted with 20 amino acids, the respective mutant libraries are predicted to have theoretical sizes of 1.6×10^5^, 3.2×10^6^ and 4.0×10^2^. These libraries were transformed into *E. coli* DH10B by electroporation, and sufficient colonies were obtained to cover the library size (measured as 7.0×10^5^, 1.5×10^8^ and 2.6×10^6^ c.f.u / mL for the three libraries, respectively). The production of phage particles was achieved through the utilization of these phagemid libraries^27, 28^. The resulting colony counts were 2.2×10^6^, 1.0×10^9^ and 2.2×10^6^ c.f.u / mL, confirming that the phage solutions had sufficient titers to evaluate the theoretical size of each library. Confirmation of the sequences of eight random clones from each library was carried out (see Fig. S2). In contrast to the TEVd sequence, the randomized residues were substituted with a range of amino acids. It was observed that the degenerate codon NNK, which includes one stop codon, was present in the Loop 1 and Loop 2 libraries. It is noteworthy that deletion mutants and frameshifts were also observed in Loop 1 and Loop 2. These may be caused by synthesis errors in the primer sequences or errors during the cloning process. Subsequently, the screening was performed on these libraries.

**Fig. 1.**
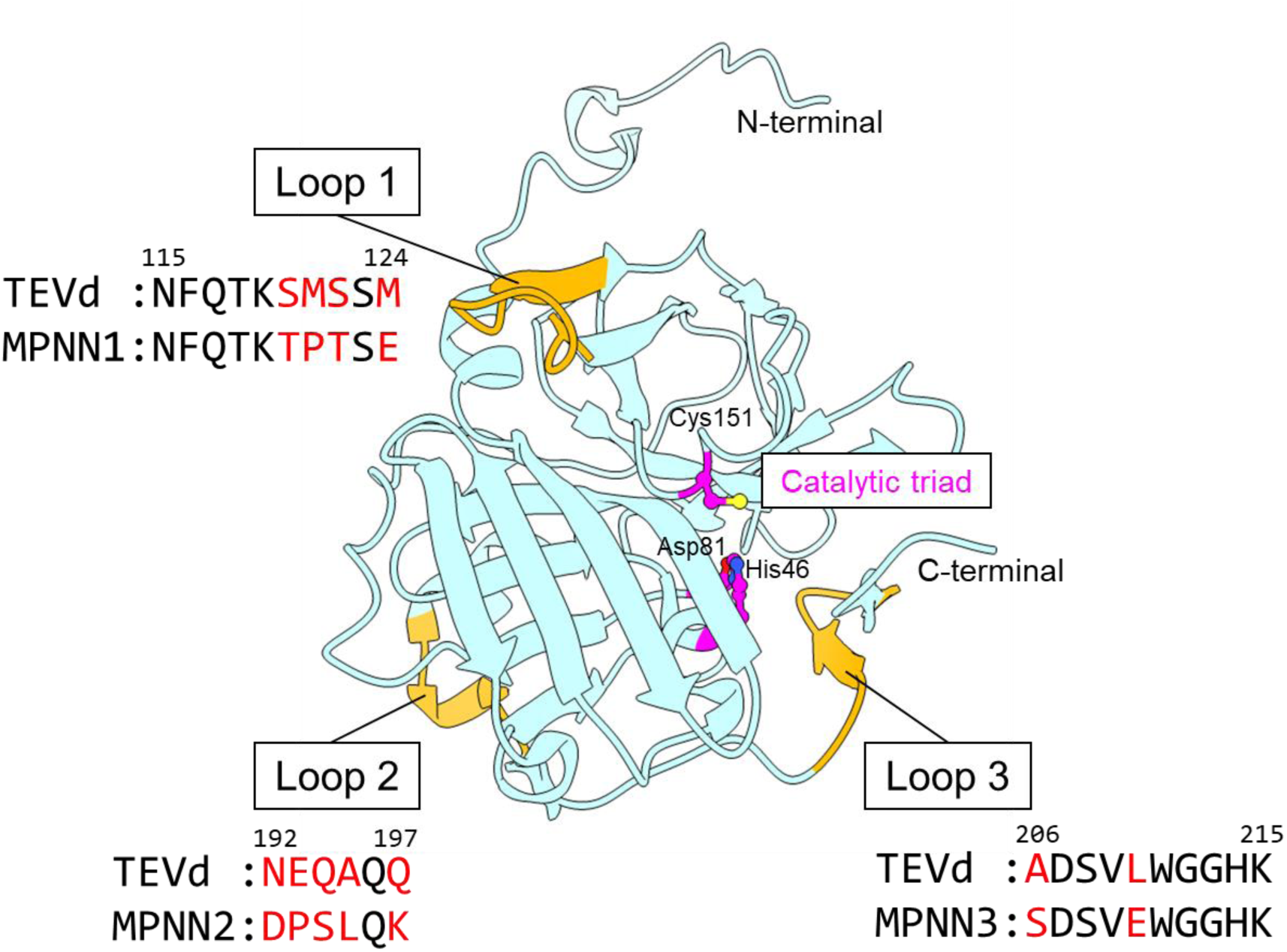
TEVd Structure showing the position of the loop regions randomized in this study. The TEVd structure^26^ (PDB: 1LVM) with the three loops designed for the library highlighted. Loops 1-3 are shown in orange and the differences in amino acid composition between the TEVd variant and the ProteinMPNN-designed HyperTEV60^15^ are shown in red. These red positions were randomised with NNK codons in the combinatorial libraries built in this study.

### Mutant screening via M13 phagemid-based system

In the present study, TEV protease mutant libraries were screened by utilizing the M13 phagemid-based selection system that was reported by A. Brödel *et al*^27, 28^. This method employs three components (Fig. 2a): a Helper Plasmid (HP), which provides the components of the M13 phage (except for the essential Genes III and VI); an Accessory Plasmid (AP), which contains a gene circuit designed to enable the conditional transcription and translation of the M13 phage Gene VI only upon TEV protease activity; and a Phagemid (PM), which carries the cloned TEV protease gene variants and Gene III. The workflow of the M13 phagemid-based selection system is illustrated in Figure 2b. Initially, a phage library is produced from the TEV protease mutant library (Step 1), which is then used to infect *E. coli* TG1 cells. During this process, the M13 phage introduces the PM into the TG1 cell (Step 2). The TEV protease variant, when transcribed and translated from the PM, is designed to recognize and cleave the TEV protease recognition site cloned into the AP, should it be active. The TEV protease recognition site is located between T7 lysozyme and T7 RNA polymerase (T7RNAP). It has been established that T7RNAP exhibits a complete loss of its transcriptional activity when T7 lysozyme is in close proximity^29^. However, upon cleavage by the TEV protease, the T7 lysozyme dissociates, and the activity of the T7RNAP is recovered^30^ (Fig. 2c). As the wild-type T7 lysozyme typically displays toxicity, a deactivated variant, C131S, is employed in this instance, through the mutation of a key active site residue^30^. It has been demonstrated that the efficiency with which the TEV protease recognition site is cleaved is directly proportional to the activity of the TEV protease gene introduced into TG1 cells by the M13 phage^30^. Consequently, a greater amount of Gene VI is transcribed by the T7 RNA polymerase for more-active TEV protease variants. Consequently, an increased number of progeny M13 phages, carrying more highly active TEV protease genes in their phagemids, are expected to be produced (Step 3). It is hypothesized that by repeating this selection cycle, high-activity TEV protease variants will thus be enriched.

**Fig. 2.**
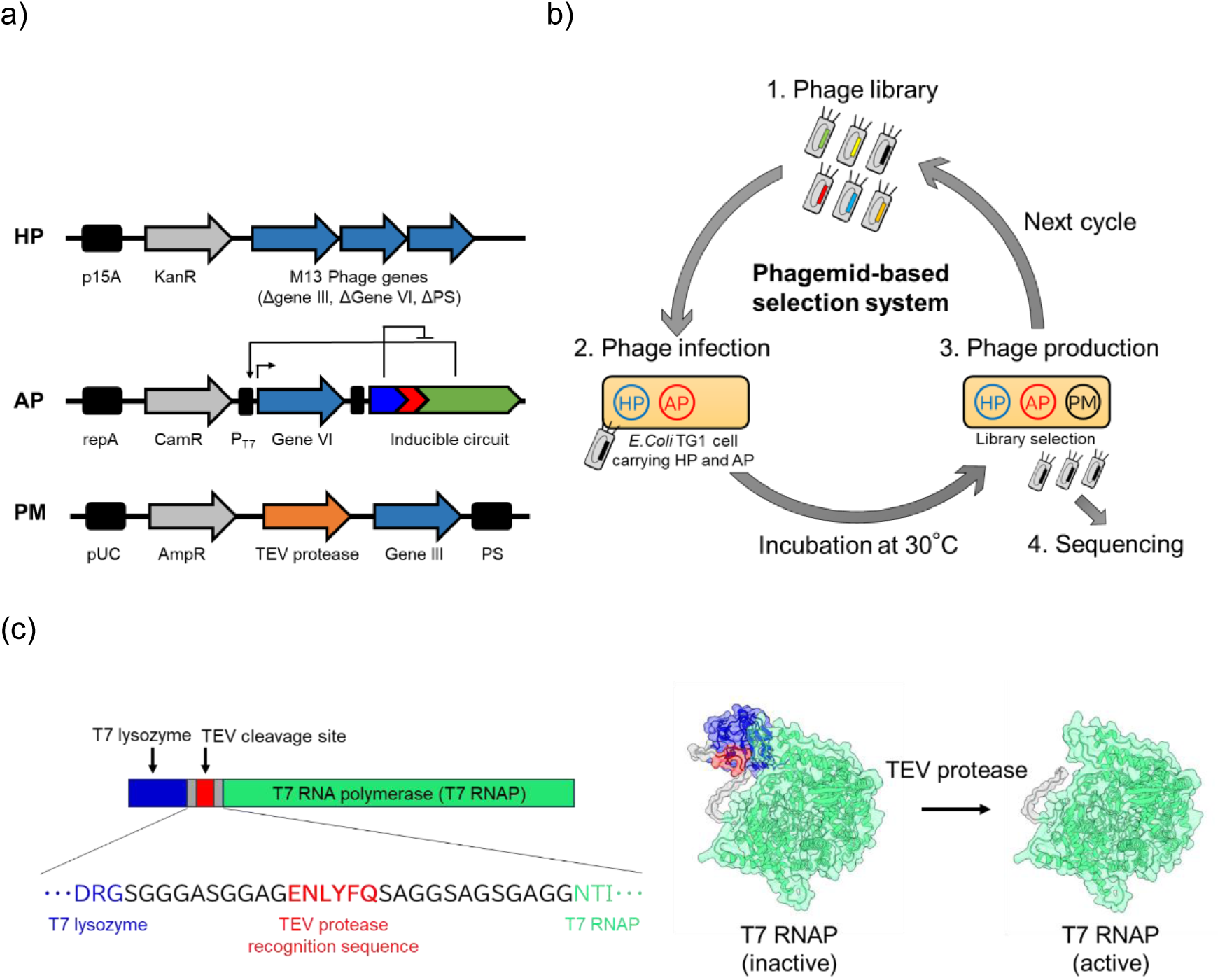
Phagemid-based selection system for TEV protease engineering. **a)** Plasmids: HP, helper phage to provide all phage genes except for gIII and gVI; AP, accessory plasmid to provide conditional Gene VI expression to enable selection of a successful evolving TEV protease variant; PM, phagemid containing an evolving TEV protease variant and gIII. **b)** Selection flow diagram: 1. Phage library: Diversification of the target gene, TEV protease, is obtained by cloning combinatorial libraries and each phage library was produced before applying the phagemid-based selection system. 2. Phage infection and 3. Phage production: Host cells, TG1, carrying the HP, AP. Only an active TEV protease activates T7 RNA polymerase and it induces Gene VI expression to complete the phage life cycle, thus enriching active library variants only. The obtained phage was either reselected in the next cycle or its phagemid sequence was confirmed by Sanger sequencing (4. Sequencing). **c)** The amino acid map and structural model of the inducible circuit. T7 RNA polymerase (T7 RNAP) was fused to the N-terminus of T7 lysozyme. The T7 lysozyme sequence used was that of an inactive mutant (C131S)^30^. A flexible linker was inserted between T7 lysozyme and T7 RNAP, containing the TEV protease recognition sequence (ENLYFQ). T7 RNAP is in an inactive state prior to cleavage by TEV protease. However, upon cleavage, T7 lysozyme dissociates, restoring transcription activity, and gene VI (encoded by the AP) is therefore conditionally transcribed. The structure model was created using AlphaFold Server^31^ (AlphaFold 3 model, Google DeepMind) and visualized using UCSF ChimeraX^32^ (version 1.7.1).

The Loop 1, Loop 2, and Loop 3 libraries were independently screened using the constructed M13 phage-based selection system. The sequences of eleven phagemids obtained after the fourth cycle for each loop are shown in Figure 3. Subsequent to the fourth cycle, the sequences for Loop 1 were analyzed, yet no particular sequence was observed to be enriched (Fig. 3a). Consequently, four additional cycles of screening were performed. Following the eighth cycle, enrichment occurred, including a variant in which six residues (commencing from position 119) were substituted with PMV-SC (with a deletion "-" at position 122). This variant was found at a frequency of 81.8%, with the remainder being the sequence KWEASY. It is noteworthy that the deletion mutant was the most enriched in Loop 1. In light of these results, the two variants for Loop 1 were prepared for further evaluation: the PMV-SC and the KWEASY variants (hereafter designated L1 and L1-1, respectively). For Loop 2, a five-residue deletion mutant was found to be highly enriched, with no other sequences detected (Fig. 3b). Consequently, this five-residue deletion mutant was prepared and designated L2. For Loop 3, three distinct variants were identified, where residues at positions 206 and 210 were substituted with SR, MA, or MP (Fig. 3c). It is evident from Figure 1 and Figure 3c that these three sequences differ from both the TEVd sequence (AL) and the ProteinMPNN sequence (SE). Furthermore, Loop 2 and Loop 3 demonstrated an apparently complete level of enrichment after four cycles. Indeed, the results of the sequencing process confirmed the absence of any alternative variants after a total of eight cycles for Loops 2 and 3. Consequently, for Loop 3, the three variants identified that substituted with MP, MA, and SR at positions 206 and 210 were prepared and designated L3, L3-1, and L3-2, respectively, for further evaluation.

**Fig. 3.**
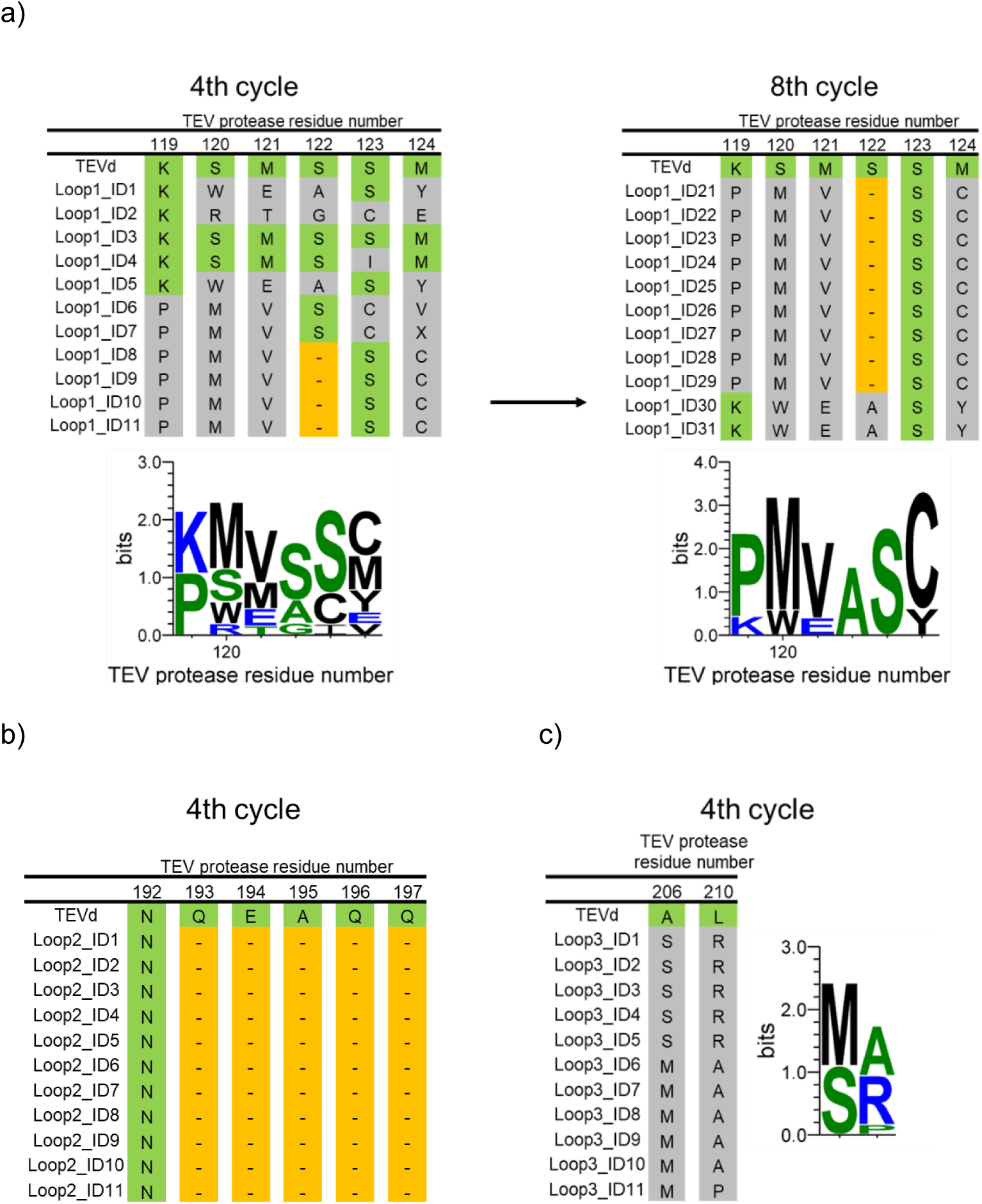
Results of screening of mutant libraries. Sequencing results of 11 selected TEV proteases. Amino acids of TEVd are highlighted in green, non-TEVd amino acids in gray, and deletion in orange. a) Sequencing results of Loop 1 after the 4^th^ and 8^th^ selection cycle, respectively. b, c) Sequencing results of Loop 2 and Loop 3 after the 4^th^ cycle, respectively. Each sequence logo was generated using Weblogo 3.7.9^33^.

### Mutant characterization

In order to investigate the effect of the individual loops designed by ProteinMPNN on TEV protease activity, variants were constructed where Loop 1, Loop 2, and Loop 3 of TEVd were substituted with the corresponding sequences from HyperTEV60 (designated MPNN1, MPNN2, and MPNN3, respectively). By evaluating the activity of these variants against a protein substrate, it was established that all three variants exhibited lower activity than TEVd (Fig. 4a). This finding indicates that the optimization of individual loops alone does not enhance TEV protease activity and, in fact, has a negative impact on the overall activity.

**Fig. 4.**
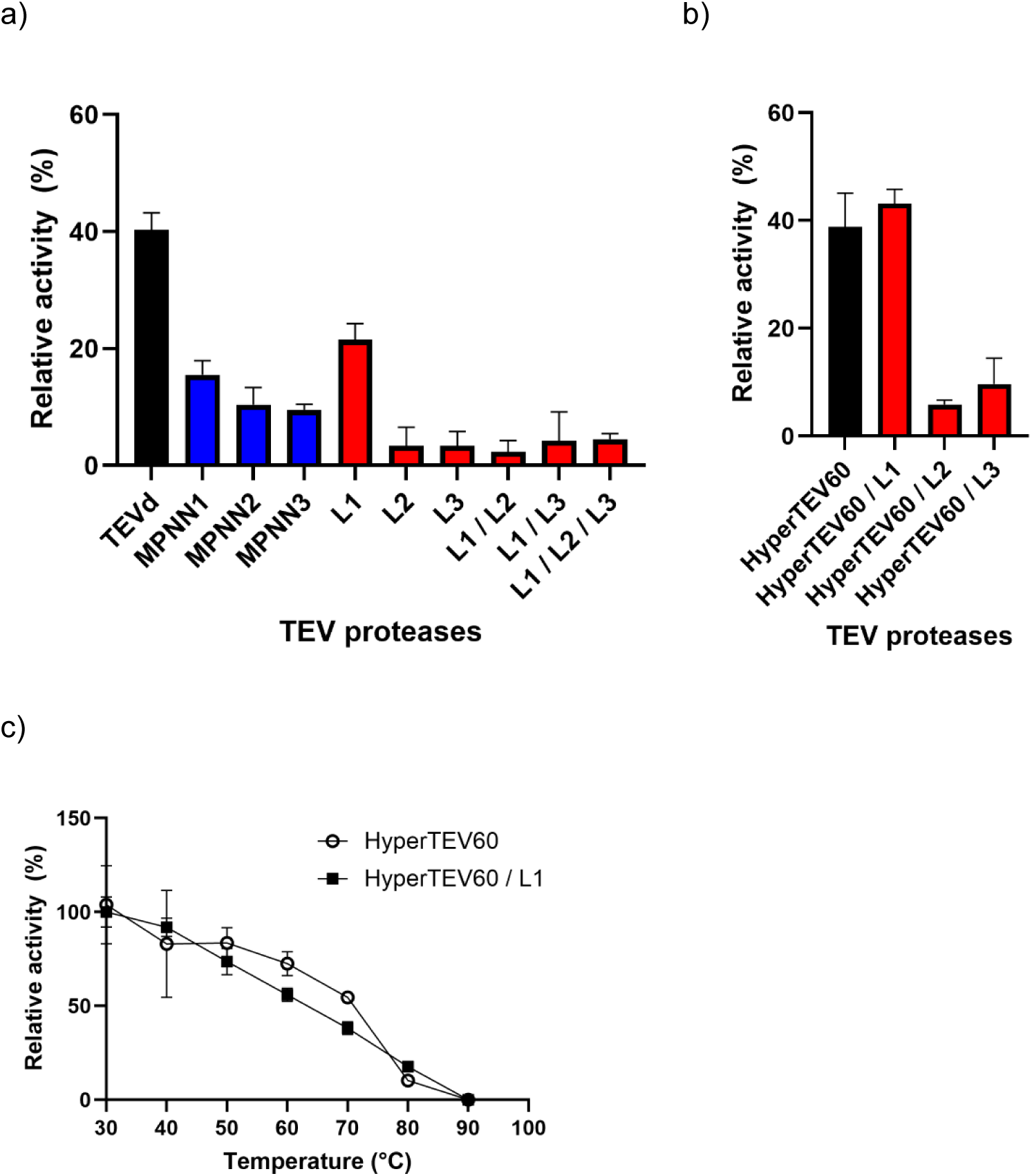
Activities and thermostabilities of TEV protease mutants. a) Relative activity of TEVd and related mutants. For each enzyme, the relative activity is expressed as the percentage of substrate consumption relative to 100% prior to the reaction. b) The activities of hyperTEV60 and selected mutants. c) Thermostability of both hyperTEV60 and hyperTEV60/L1. Here, the activity after 10 minutes of heat treatment at each temperature is shown as relative activity compared to the activity at 30°C. Data are mean ± s.e.m. from three independent experiments (*n* = 3).

Subsequently, the six variants obtained from the M13 phage-based selection system were expressed and purified in *E. coli*. However, the expression levels of L1-1, L3-1, and L3-2 were found to be significantly reduced, which made it extremely difficult to obtain sufficient quantities for activity measurements. Consequently, a comparison was made between the activities of the purified variants L1, L2, and L3 and TEVd (Fig. 4a). Among these variants, L1 exhibited the highest activity, although both L2 and L3 showed significantly reduced activity. Despite the elevated activity exhibited by L1 in comparison to the three MPNN variants (MPNN1, 2, and 3), its activity remained lower than that of TEVd. Furthermore, the multiple mutants combining L1, L2, and L3 also exhibited significantly reduced activity in comparison to L1 (Fig. 4a). The rationale behind the acquisition of variants exhibiting reduced activity relative to TEVd, despite the theoretical presence of the TEVd sequence within the mutant libraries, is likely attributable to the preferential production of M13 phage by the mutants, attributable to their apparent heightened cleavage efficiency under selection conditions, as indicated by expression level or other system components.

An investigation was then conducted into the effect of L1, L2, and L3 on the hyperTEV60 scaffold. To this end, three variants were created, in which each respective loop of HyperTEV60 was substituted with the selected sequence. The activities of these variants were then evaluated. It is noteworthy that the HyperTEV60/L1 variant exhibited activity that was almost equivalent to that of the parental HyperTEV60 (Fig. 4b). In contrast, L2 and L3 were found to reduce the activity of the HyperTEV60 scaffold as well. A comparison of the kinetic parameters of HyperTEV60 and HyperTEV60/L1 revealed that the *K*_m_ of HyperTEV60/L1 was reduced to approximately one-third that of HyperTEV60, suggesting an improvement in substrate affinity (see Table 1 and Fig. S3). Despite the *k*_cat_ of HyperTEV60 being reduced to 43.9% due to the L1 modification, the resulting *k*_cat_/*K*_m_ of HyperTEV60/L1 was very similar to that of HyperTEV60 (and even marginally higher, although not significantly so) This finding indicates that engineering Loop 1 has a significant impact on enhancing the *K*_m_ of TEV protease while maintaining the *k*_cat_/*K*_m_. It is hypothesized that this effect is dominant within the M13 phage-based selection system, where the intracellular detection of TEV protease relies on a low concentration of the substrate protein, rendering *K*_m_ the controlling factor. In contrast, the deep-learning-based ProteinMPNN was previously shown to enhance the *k*_cat_/*K*_m_ of TEVd primarily by improving the *k* ^15^. This finding demonstrates that the directed evolution approach used in this study and the deep-learning-based tool exhibit different search tendencies in the optimization landscape. With regard to thermal stability, both enzymes demonstrated a significant loss of activity at 80°C, with almost complete inactivation occurring after 10 minutes of treatment at 90°C. While HyperTEV60 exhibited marginally higher thermal stability within the temperature range of 50°C to 70°C, it was determined that the modification of Loop 1 did not result in a substantial alteration to the overall thermal stability. This finding indicates that, in addition to the loop, other regions of the TEV protease play a pivotal role in its folding process.

**Table 1.** Kinetic parameters of HyperTEV60 and HyperTEV60 / L1.

|  | HyperTEV60 | HyperTEV60/L1 |
| --- | --- | --- |
| $K_m$ ($\mu\text{M}$) | 18.61 | 6.38 |
| $k_{\text{cat}}$ ($\text{min}^{-1}$) | 1.32 | 0.58 |
| $k_{\text{cat}} / K_m$ ($\mu\text{M}^{-1} \cdot \text{min}^{-1}$) | 0.07 | 0.09 |

An investigation was conducted to ascertain the rationale behind the *K*_m_ enhancement observed in HyperTEV60/L1. This investigation entailed the detailed examination of structural models of HyperTEV60 and HyperTEV60/L1, which were constructed utilizing the Boltz-2 approach^34^. Thus, we constructed substrate peptide complex structures that were analogous to the crystal structure^26^ (PDB ID: 1LVM). The root-mean-square deviation (RMSD) of the Cα atoms relative to the crystal structure was 0.332Å for HyperTEV60 and 0.303Å for HyperTEV60/L1, confirming the substantial consistency of the substrate peptide structures (Fig. S4). This finding suggests that the models have been adequately predicted, thus paving the way for further structural analyses. A comparison of the two structures revealed a discrepancy in the conformation of Loop 1 (highlighted in yellow in Fig. 5). Specifically, Loop 1 in HyperTEV60/L1 exhibited a structure that moved away from the substrate peptide binding region in comparison to HyperTEV60, resulting in an expansion of the gap in the substrate binding pocket. This finding indicates that the modified Loop 1 may facilitate the binding of the substrate protein, which is hypothesized to contribute to the observed *K*_m_ improvement in HyperTEV60/L1.

**Fig. 5.**
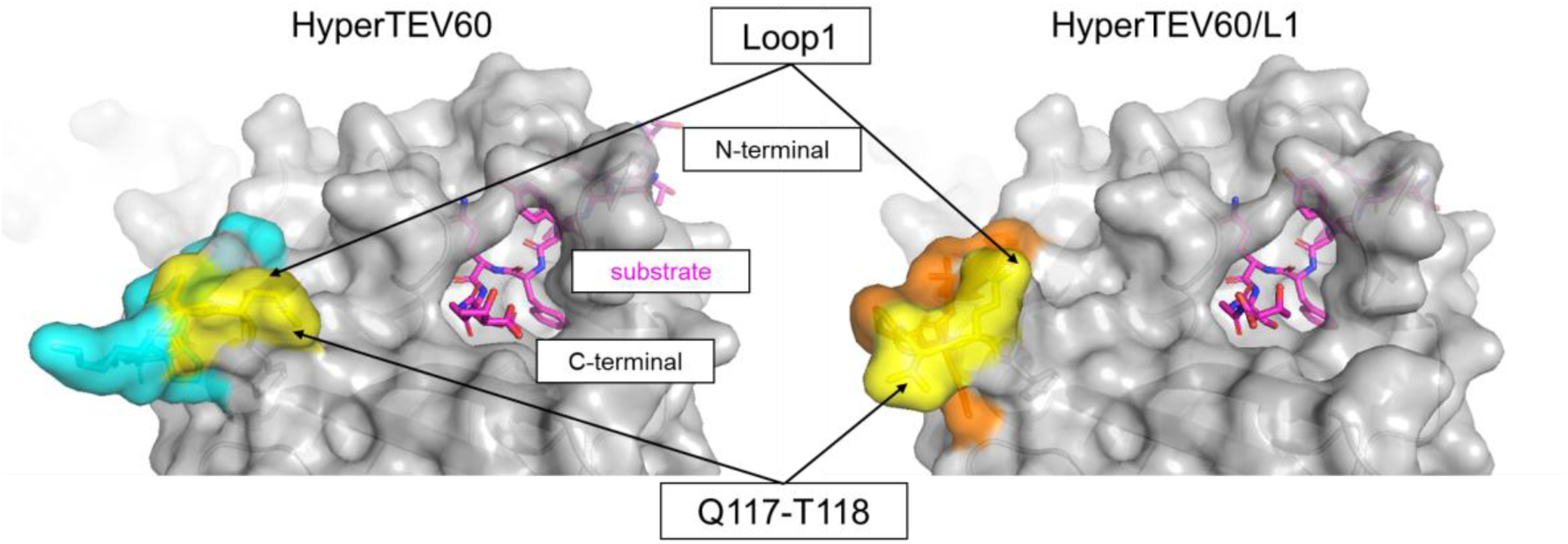
Structural model of hyperTEV60 and hyperTEV60/L1 with substrate. Each structural model was predicted using Boltz-2^34^, with the full-length amino acid sequences of hyperTEV60 and hyperTEV60/L1 as queries, to generate complex models with the substrate peptide (TTENLYFQSGT). The substrate peptide sequence used for this model was identical to that used for the crystallization of the TEV protease complex^26^ (PDB ID: 1LVM). Each structure is depicted as a surface model (grey), with the substrate peptide shown as stick model in magenta. The two residues Q117-T118 are highlighted in yellow, while Loop1 is represented by HyperTEV60 (cyan) and HyperTEV60/L1 (orange) respectively. The structure models were visualized using PyMOL (version 3.4.1.4).

## Conclusion

In this study, we aimed to answer the important question of what types of variants are produced by directed evolution versus deep-learning-based design methods within the vast protein space for enzyme loop engineering. To achieve this, we constructed mutant libraries for three loops of TEV protease and performed screening using the M13 phagemid-based selection system. As a result, we identified a Loop 1 sequence that significantly improves the *K*_m_ of HyperTEV60, which is one of the variants with the highest activity reported to date. The Loop 1 sequence discovered through experimental evolution includes a single-residue deletion. This work demonstrates the utility of the M13 phagemid-based selection system for enzyme loop engineering and refocuses attention on the importance of evaluating deletion mutants. The improved *K*_m_ is presumed to result from the expansion of the substrate binding pocket due to the Loop 1 mutation and deletion. This suggests that Loop 1 plays a critical role in substrate affinity in TEV protease. However, as the *k*_cat_ was reduced, the resulting *k*_cat_/*K*_m_ of HyperTEV60/L1 was very similar to that of HyperTEV60 (marginally higher, although not significantly so). By combining deep-learning-based tools for initial global optimization and directed evolution methods for localized optimization, we demonstrate the potential to obtain kinetically-tuned variants.

## Materials and methods

### Strains and media

Standard DNA cloning was performed with chemically competent DH10B (Thermo Fisher Scientific) and TG1 cells (Zymo Research). Combinatorial library cloning was performed with DH10B electrocompetent cells or electrocompetent TG1 cells. Selection phage production was carried out with DH10B cells. All phage-assisted selection experiments were performed with TG1 cells. The expression of TEV mutant was performed with BL21 (DE3) pLys cells. Genotypes of all strains are listed in Supplementary Table. Cells were grown in Luria-Bertani medium (Sigma-Aldrich), M9 minimal medium (6.8 g / L Na_2_HPO_4_, 3.0 g / L KH_2_PO_4_, 0.5 g / L NaCl, 1.0 g / L NH_4_Cl, 2 mM MgSO_4_, 100 mM CaCl_2_, 0.2% (w/v) glucose, 1 mM thiamine-HCl), 2 × TY medium (5 g / L NaCl, 10 g / L yeast extract, 16 g / L tryptone) or S.O.C. medium (Sigma-Aldrich). Chloramphenicol (25 μg / ml), kanamycin (50 μg / ml) and carbenicillin (100 μg / ml) were added where appropriate. *E. coli* strains used in this research are shown in Table S1.

### Cloning and plasmid construction

The sequences for HP, AP, and PM were obtained from Addgene or from plasmids held by our laboratory, as described in prior studies^30^. Specifically, the M13KO7 helper phage plasmid (HP), which has the weak M13 packaging signal (PS), gene III, and gene VI removed, was obtained from Addgene plasmid ID 80852. Next, the accessory plasmid (AP) was constructed using Addgene plasmid ID 134356 as a template, referencing prior literature^35^, to enable transcription and translation of Gene VI via TEV protease activity. The 1155–1926 region of the template plasmid was removed, and instead, a gene fusing T7 Lysozyme to T7 RNA polymerase via a TEV cleavage site was cloned (Fig. S5). Since T7 Lysozyme exhibits cytotoxicity, the C131S mutant was used, which is inactivated by introducing a mutation into the active site^30^. The promoter for this fusion gene utilized the pKAT promoter^36^. Finally, for PM, the Addgene plasmid ID 80852 was used as a template, and the gene for TEV protease (GenBank no. NC_001555.1) was cloned in place of transcription factor cl (Fig. S6). To suppress self-cleavage, the S219D mutant was used as the wild-type TEV protease^26^. To suppress the leaky expression of the TEV protease gene, the araC regulatory gene was introduced upstream of the TEV protease coding sequence, and the promoter used was pBAD. Plasmids were purified using the QIAprep Spin Miniprep Kit or the HiSpeed Plasmid Maxi Kit (QIAGEN). Nucleotide sequences of all cloned constructs were confirmed by DNA sequencing (Eurofins or FCL). The each amino acids sequences of genes are listed in Fig. S7.

### Construction of combinatorial TEV protease libraries

Combinatorial libraries were cloned based on forward and reverse primers containing NNK codons (where N = A, C, G or T and K = G, T) at the randomized positions (Table S2). Using these primers, the PM (for cloning the wild-type TEV protease) was linearized via inverse PCR. Fragments obtained by gel extraction after agarose gel electrophoresis were ligated using KLD Enzyme Mix (NEB). Randomized positions of TEV protease were as follows: Library 1 (S120, M121, S122, M124); Library 2 (N192, Q193, E194, A195, Q197); Library 3 (A206, L210). Cells were transformed and plated on 24 cm^2^ Nunc BioAssay Dishes (Thermo Fisher Scientific). Transformation efficiency was estimated by colony counting of plated serial dilutions. The next day, colonies were harvested and phagemid DNA was purified. All combinatorial libraries and corresponding sequencing results are listed in Fig. S2.

### Selection phage production

Selection phage production was performed in DH10B cells containing HP: M13KO7-ΔPS-ΔgeneIII (Addgene plasmid ID 134351). Cells were made electrocompetent, phagemids transformed and cells were grown overnight at 30°C in 2 × TY medium supplemented with 50 μg/mL kanamycin and 100 μg/mL ampicillin. Cell cultures were centrifuged for 10 min at 4,000 × g and supernatants were sterile filtered (0.22 μm pore size, Millex-GV). The phage titer was determined by infecting TG1 cells with phage diluted in 2 × YT medium. This was achieved by mixing 50 μL of the diluted phage solution with 450 μL of TG1 cells at an OD of 0.4 to 0.6. After culturing at 37°C for 1 hour at 200 rpm, the mixture was inoculated onto LB plates containing 100 μg/mL ampicillin. Plates were incubated overnight at 37°C, and colonies were counted the following morning for analysis.

### Phage-assisted selections

TG1 cells containing the HP and the AP were grown on M9 minimal medium plates supplemented with 50 μg/mL chloramphenicol and 50 μg/mL kanamycin. Starter cultures were grown for 6-8 h until the OD_600_ reached 0.4–0.8. 5–10 ml of the starter cultures was infected with selection phages at a MOI of 0.1–10. Samples were kept at 37°C without stirring for 10 min before they were incubated for 18–20 h at 30°C and 200 r.p.m (Stuart Shaking Incubator SI500). Overnight cultures were centrifuged for 10 min at 4,000 × g and the phage supernatant was used to start a new round of selection. After four to eight rounds of selection, phage supernatants were sterile filtered and diluted, before infecting TG1 cells. Infected cells were inoculated on ampicillin plates and eleven colonies per selection were grown overnight in LB medium. Phagemid DNA was purified using the QIAprep Spin Miniprep Kit (QIAGEN) and analyzed by Sanger sequencing.

### Expression and Purification of each mutant

The pET-26b plasmid in which each TEV protease was cloned was constructed by Gibson assembly. TEV protease was fused with the His-tag sequence at the N-terminus. The plasmid was introduced into *E. coli* BL21 (DE3) pLys, and colonies were inoculated in LB medium containing 100 μg/ml ampicillin at 37°C. His-tag fusion proteins were expressed by induction using 1 mM isopropyl-β-D-thiogalactopyranoside when the OD_600_ reached 0.6–1.0. The temperature was decreased to 25°C, followed by incubation for 16 h. Harvested cells were washed with saline, then re-suspended in B-PER containing DNase I and Lysozyme (Thermo Fisher Scientific). After incubation for 15 min at room temperature, the lysate was centrifuged for 5 min at 15,000 × g. The supernatant was added to a His SpinTrap TALON column (Cytiva), and bound proteins were washed with a wash buffer (50 mM Tris-HCl, 300 mM NaCl, 50 mM imidazole, pH 8.0) and eluted with an elution buffer (50 mM Tris-HCl, 300 mM NaCl, 250 mM imidazole, pH 8.0). The eluted fraction buffer was exchanged for a solution of 50 mM Tris-HCl and 300 mM NaCl at pH 8.0 using an Amicon 3K (Merck), after which the sample was immediately subjected to an enzymatic assay.

### Enzymatic assay

The activity measurement of wild-type and mutant TEV proteases was performed according to previous studies^30^. Specifically, 5 μL of TEV protease prepared at 0.01 mg/mL was mixed with 5 μL of substrate protein at 2 mg/mL and reacted for 3 hours at 30°C for screening of mutants. The substrate protein used was the MBP-TEV cleavage site-FKBP-EGFP fusion protein described in the literature^15^. The amino acid sequence of the substrate protein, as reported in the literature, was chemically synthesized (Twist Bioscience) and cloned into pET-26b using the HiFi DNA Assembly Kit (NEB). Expression and purification were performed using the same methods as for the TEV protease. The reaction was terminated by adding 10 μL of 2 × SDS-PAGE sample buffer (Invitrogen) and heating at 95°C. Subsequently, the extent of substrate cleavage from the SDS-PAGE bands was analyzed using ImageJ. Enzymatic parameters were obtained using the following method based on a previous study^37^. Reactions were carried out with 0.05–0.1 μM protease and 50 μM to 2 mM substrate for evaluation of kinetic parameters. Proteases and substrates were prepared as solutions at 2 × concentration (5 µl each) then combined to yield a total reaction volume of 10 µl. Reactions were incubated at 30 °C for 20 min and quenched with 10 µl of 2 × SDS-PAGE sample buffer. Prior to conducting reactions in triplicate, all conditions were tested and monitored at 5, 10, 30 min to ensure that 20 min was within the linear range of the reaction (<25% substrate consumption). The cutting efficiency was calculated from the area values of the bands before and after cutting, and fit to the Lineweaver-Burk equation so as to calculate *K*_m_ and *k*_cat_.

## Data availability

Sequences and plasmid maps (as listed in Fig. S5 and S6), as well as DNA plasmids have been deposited and are available from Addgene: https://www.addgene.org/. All other relevant data are available from the corresponding author on request.

## Acknowledgement

This work was supported by Ajinomoto Co., Inc. We would like to thank Amir Guppy, Dario Bazzoli, Qi Lin, Olga Komarynets, Shayma Bukannan and Haobin Chen for their support. We would also like to thank Andreas K. Brödel for his valuable advice.

## Author contributions

H. Y.: conceptualization, methodology, mutant design, screening, mutant characterization and writing. M. I.: supervision, conceptualization, methodology and writing. All authors reviewed the manuscript.

