## Supplementary_information for "Loop engineering and activity improvement of TEV protease by a phagemid-based selection system"

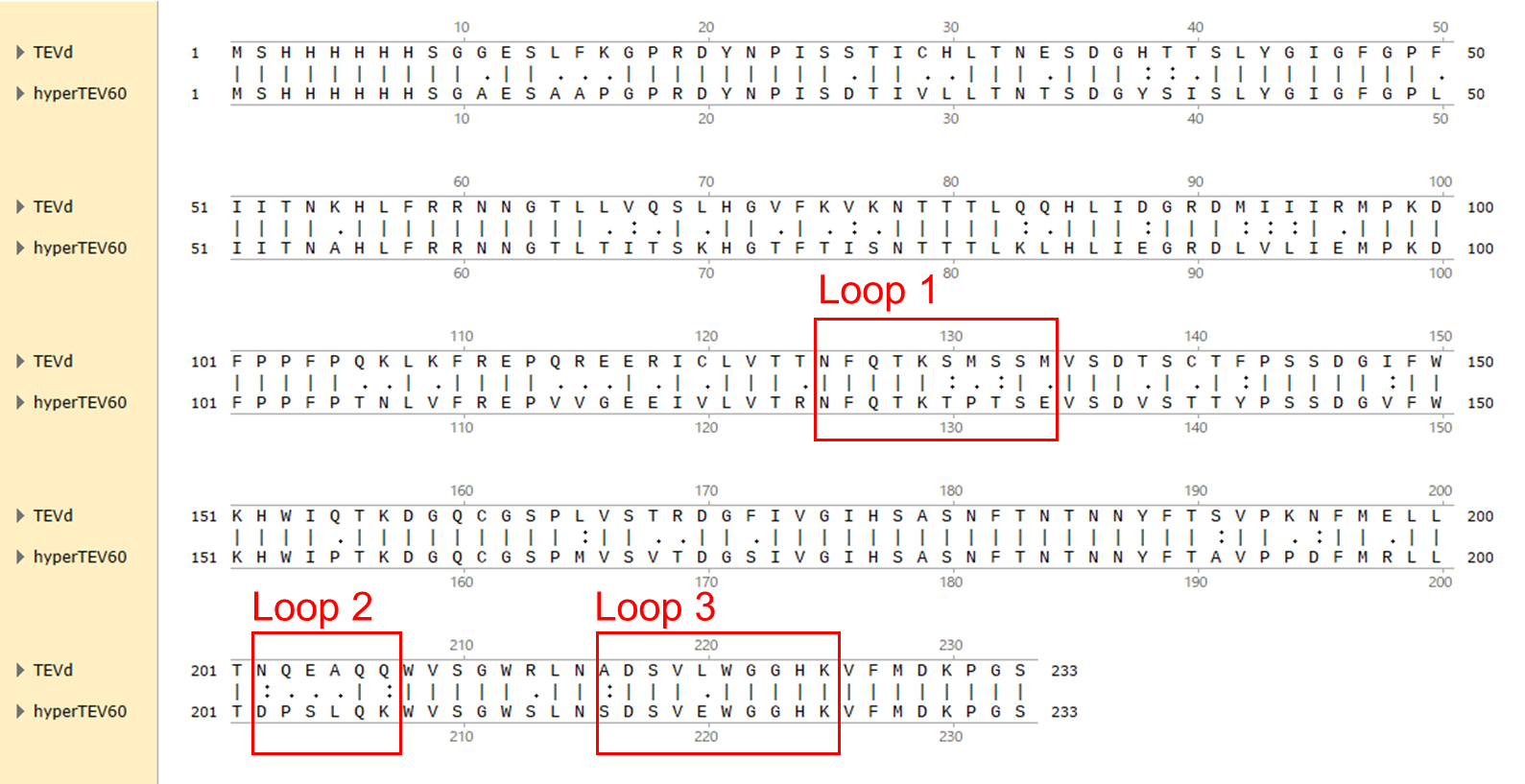

**Supplementary Fig. 1 Sequence alignment between TEVd and hyperTEV60**

TEVd and hyperTEV60 sequence alignments were performed using the software SnapGene version 8.0.2.

a)	b)

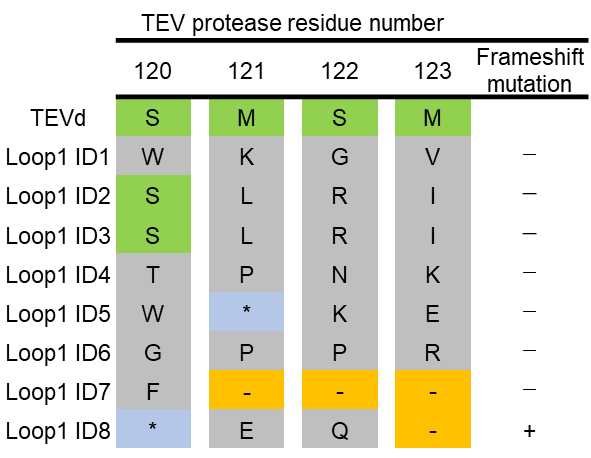

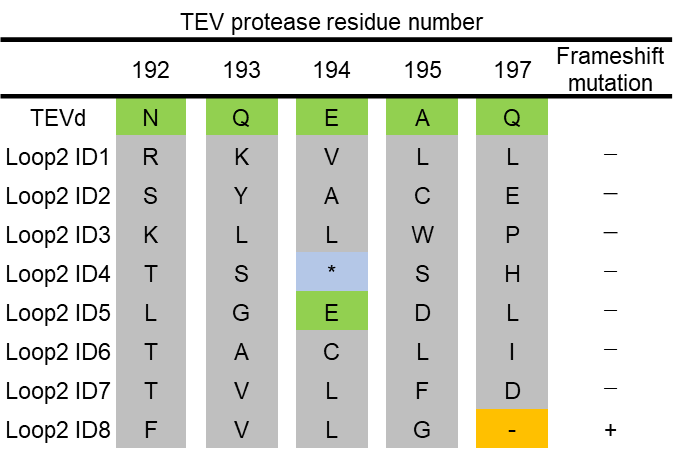

c)

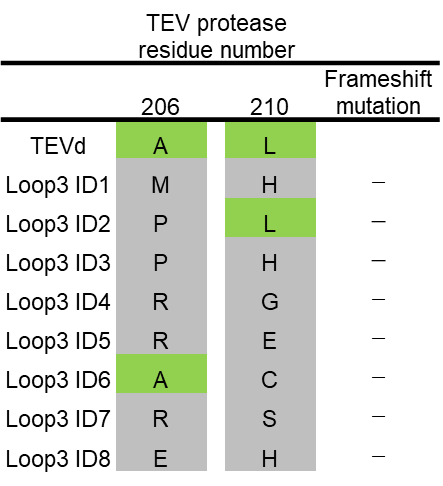

**Supplementary Fig. 2 Sequencing results of the three libraries before screening**

Sequencing results of 8 selected TEV proteases in a) Loop 1, b) Loop 2, and c) Loop 3. Amino acids of TEVd are highlighted in green, non-TEVd amino acids in gray, stop codon in light blue and deletion in orange. Mutants in which a frameshift occurred due to deletion of residues are denoted with “+”.

a)

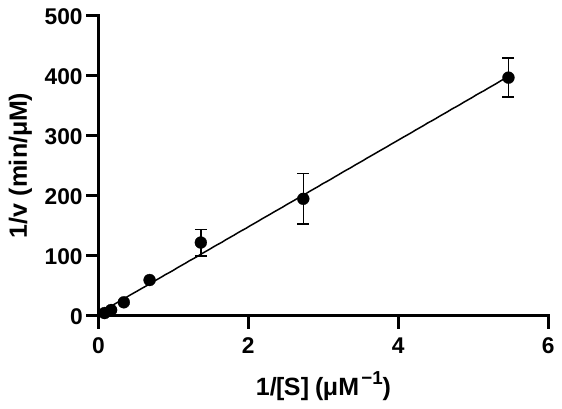

b)

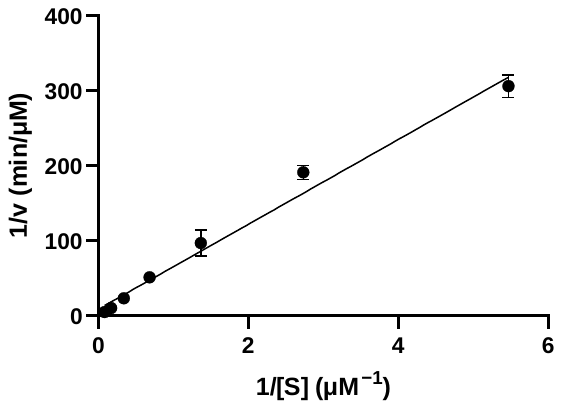

**Supplementary Fig. 3 Lineweaver-Burk plot of hyperTEV60 and hyperTEV60/L1**

a) Lineweaver-Burk plot of hyperTEV60. b) hyperTEV60/L1. Data are mean ± s.e.m. from three independent experiments (*n* = 3).

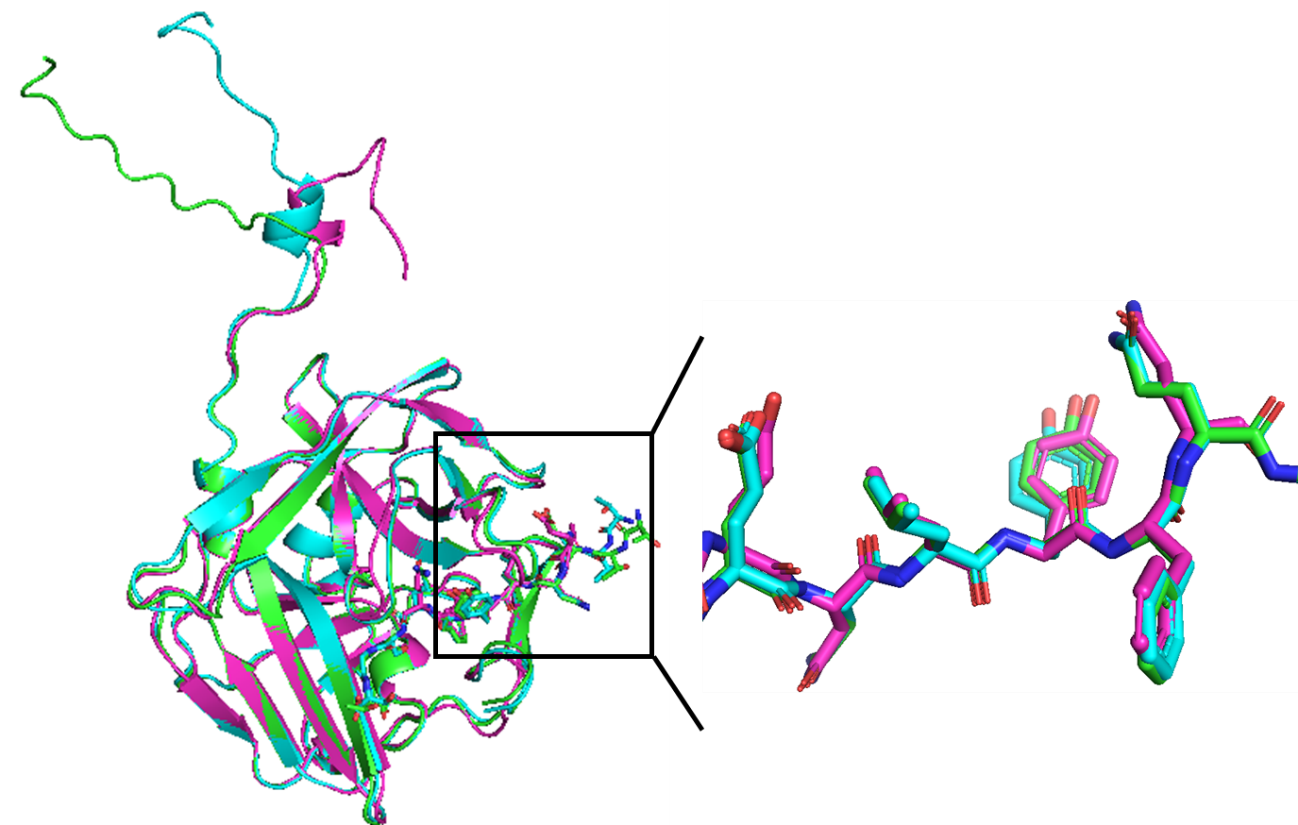

**Supplementary Fig. 4 Structural comparison among TEV proteases**

Comparison of the crystal structure of TEVd and substrate peptide (magenta, PDB ID: 1LVM) with the predicted complex structures of HyperTEV60 (green) and HyperTEV60/L1 (cyan) generated by Boltz-2. The structures of the three TEV proteases are shown as cartoon models, while the substrate peptides are displayed as stick models. An enlarged view of the substrate peptide is shown on the right. Structure alignment and rendering were performed using PyMOL (version 3.4.1.4).

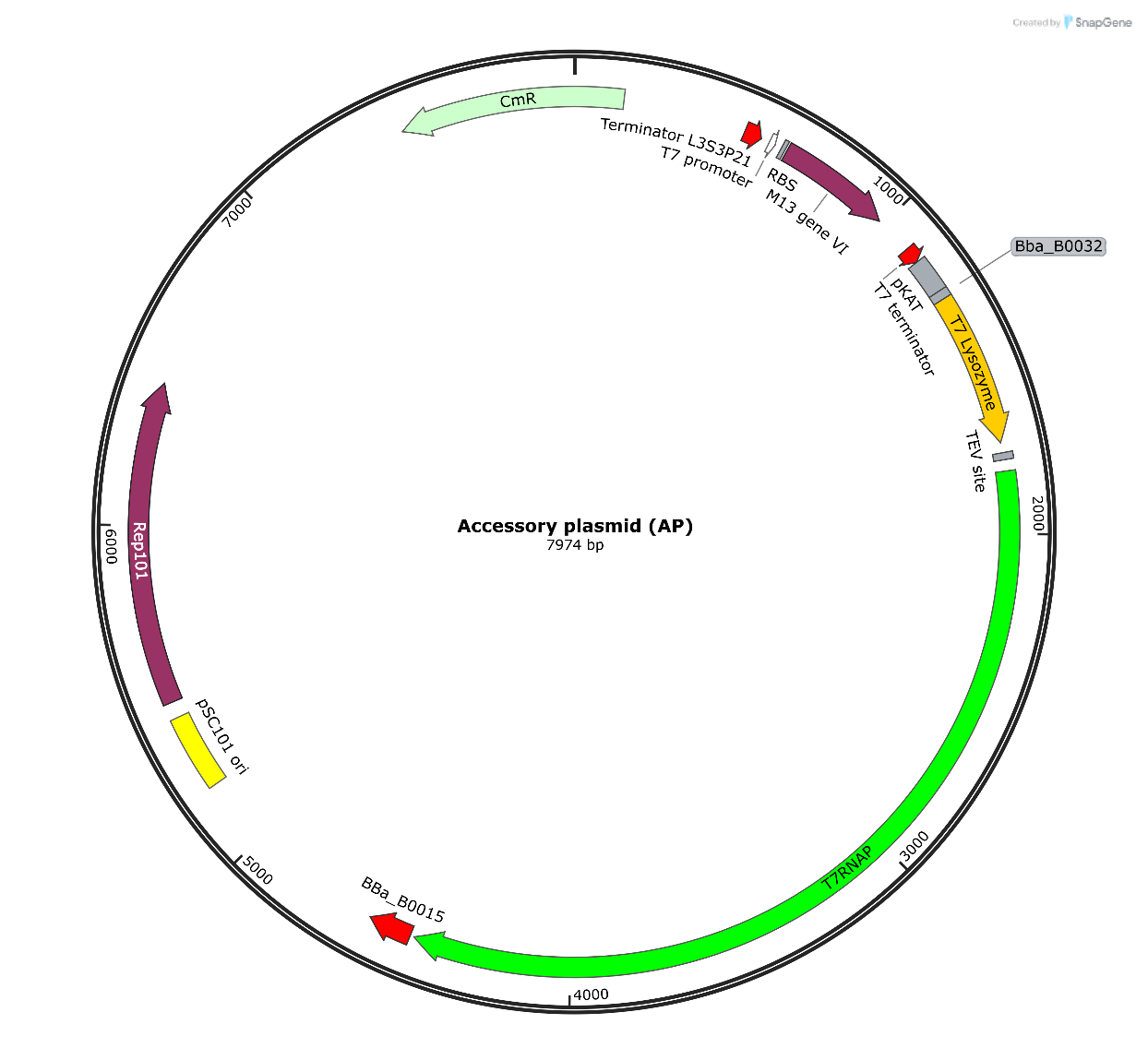

**Supplementary Fig.5 Vector map of accessory plasmid (AP) used in this study.**

This vector map was drawn using the software SnapGene version 8.0.2.

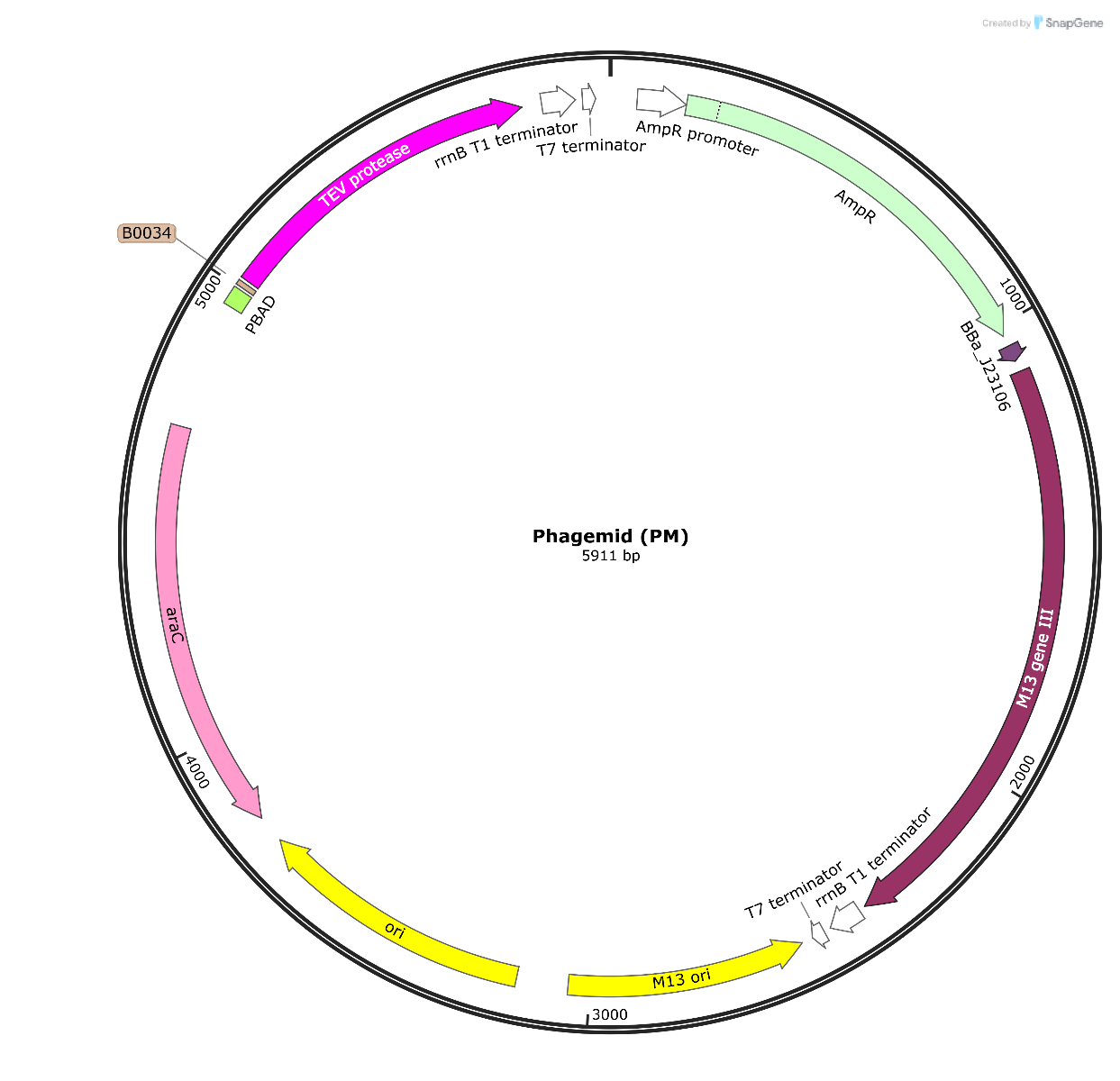

**Supplementary Fig.6 Vector map of phagemid (MP) used in this study.**

This vector map was drawn using the software SnapGene version 8.0.2.

>TEVd (S219D)

MSHHHHHHSGGESLFKGPRDYNPISSTICHLTNESDGHTTSLYGIGFGPFIITNKHLFRRNNGTLLVQSLHGVFKVKNTTTLQQHLIDGRDMIIIRMPKDFPPFPQKLKFREPQREERICLVTTNFQTKSMSSMVSDTSCTFPSSDGIFWKHWIQTKDGQCGSPLVSTRDGFIVGIHSASNFTNTNNYFTSVPKNFMELLTNQEAQQWVSGWRLNADSVLWGGHKVFMDKPGS

>HyperTEV60

MSHHHHHHSGAESAAPGPRDYNPISDTIVLLTNTSDGYSISLYGIGFGPLIITNAHLFRRNNGTLTITSKHGTFTISNTTTLKLHLIEGRDLVLIEMPKDFPPFPTNLVFREPVVGEEIVLVTRNFQTKTPTSEVSDVSTTYPSSDGVFWKHWIPTKDGQCGSPMVSVTDGSIVGIHSASNFTNTNNYFTAVPPDFMRLLTDPSLQKWVSGWSLNSDSVEWGGHKVFMDKPGS

>HyperTEV60/L1

MSHHHHHHSGAESAAPGPRDYNPISDTIVLLTNTSDGYSISLYGIGFGPLIITNAHLFRRNNGTLTITSKHGTFTISNTTTLKLHLIEGRDLVLIEMPKDFPPFPTNLVFREPVVGEEIVLVTRNFQTPMVSCVSDVSTTYPSSDGVFWKHWIPTKDGQCGSPMVSVTDGSIVGIHSASNFTNTNNYFTAVPPDFMRLLTDPSLQKWVSGWSLNSDSVEWGGHKVFMDKPGS

>T7Lyso-TEVcs-T7RNAP

MARVQFKQRESTDAIFVHCSATKPSQNVGVREIRQWHKEQGWLDVGYHFIIKRDGTVEAGRDEMAVGSHAKGYNHNSIGVCLVGGIDDKGKFDANFTPAQMQSLRSLLVTLLAKYEGAVLRAHHEVAPKASPSFDLKRWWEKNELVTSDRGSGGGASGGAGENLYFQSAGGSAGSGAGGNTINIAKNDFSDIELAAIPFNTLADHYGERLAREQLALEHESYEMGEARFRKMFERQLKAGEVADNAAAKPLITTLLPKMIARINDWFEEVKAKRGKRPTAFQFLQEIKPEAVAYITIKTTLACLTSADNTTVQAVASAIGRAIEDEARFGRIRDLEAKHFKKNVEEQLNKRVGHVYKKAFMQVVEADMLSKGLLGGEAWSSWHKEDSIHVGVRCIEMLIESTGMVSLHRQNAGVVGQDSETIELAPEYAEAIATRAGALAGISPMFQPCVVPPKPWTGITGGGYWANGRRPLALVRTHSKKALMRYEDVYMPEVYKAINIAQNTAWKINKKVLAVANVITKWKHCPVEDIPAIEREELPMKPEDIDMNPEALTAWKRAAAAVYRKDKARKSRRISLEFMLEQANKFANHKAIWFPYNMDWRGRVYAVSMFNPQGNDMTKGLLTLAKGKPIGKEGYYWLKIHGANCAGVDKVPFPERIKFIEENHENIMACAKSPLENTWWAEQDSPFCFLAFCFEYAGVQHHGLSYNCSLPLAFDGSCSGIQHFSAMLRDEVGGRAVNLLPSETVQDIYGIVAKKVNEILQADAINGTDNEVVTVTDENTGEISEKVKLGTKALAGQWLAYGVTRSVTKRSVMTLAYGSKEFGFRQQVLEDTIQPAIDSGKGLMFTQPNQAAGYMAKLIWESVSVTVVAAVEAMNWLKSAAKLLAAEVKDKKTGEILRKRCAVHWVTPDGFPVWQEYKKPIQTRLNLMFLGQFRLQPTINTNKDSEIDAHKQESGIAPNFVHSQDGSHLRKTVVWAHEKYGIESFALIHDSFGTIPADAANLFKAVRETMVDTYESCDVLADFYDQFADQLHESQLDKMPALPAKGNLNLRDILESDFAFA

**Supplementary Fig.7 Amino acid sequences**

**Supplementary Table 1 *E. coli* strains used in this research**

| Stain | Genotype | Company |
| --- | --- | --- |
| DH10B | F– *mcrA* Δ(*mrr-hsdRMS-mcrBC*) φ80*lacZ*ΔM15 Δ*lacX74 recA1 endA1 araD139* Δ (*ara-leu*)7697 *galU galK* λ– *rpsL*(Str^R^) *nupG* | Thermo Fisher Scientific |
| TG1 | F’[traD36 lacIq ∆(lacZ) M15 proA+B+] glnV (supE) thi-1 ∆(mcrB-hsdSM)5 (rK- mK- McrB-) thi ∆(lac-proAB) | Zymo Research |
| BL21 (DE3) pLys | F^–^*omp*T *hsd*SB (r_B_–, m_B_–) *gal dcm* (DE3) pLysS(CamR) | Thermo Fisher Scientific |

**Supplementary Table 2 Primers for library construction**

| No. | name | sequence |
| --- | --- | --- |
| 1 | Library_1_FW | [Pho]ANNKNNKNNKAGCNNKGTGTCGGATACCTCTTGTAC |
| 2 | Library_1_RV | [Pho]TTCGTCTGAAAGTTCGTGGTAACC |
| 3 | Library_2_FW | [Pho]CNNKNNKNNKNNKCAGNNKTGGGTTTCAGGTTGGCGTCTTAATG |
| 4 | Library_2_RV | [Pho]GTCAGCAGTTCCATGAAGTTCTTGGGG |
| 5 | Library_3_FW | [Pho]TNNKGATTCGGTANNKTGGGGTGGTCACAAAGTGTTCATG |
| 6 | Library_3_RV | [Pho]TTAAGACGCCAACCTGAAACCCATTGCTG |
